# Thiocyanate and organic carbon inputs drive convergent selection for specific autotrophic *Afipia* and *Thiobacillus* strains within complex microbiomes

**DOI:** 10.1101/2020.04.29.067207

**Authors:** Robert J. Huddy, Rohan Sachdeva, Fadzai Kadzinga, Rose Kantor, Susan T.L. Harrison, Jillian F. Banfield

## Abstract

Thiocyanate (SCN^-^) contamination threatens aquatic ecosystems and pollutes vital fresh water supplies. SCN^-^ degrading microbial consortia are commercially deployed for remediation, but the impact of organic amendments on selection within SCN^-^ degrading microbial communities has not been investigated. Here, we tested whether specific strains capable of degrading SCN^-^ could be reproducibly selected for based on SCN^-^ loading and the presence or absence of added organic carbon. Complex microbial communities derived from those used to treat SCN^-^ contaminated water were exposed to systematically increased input SCN concentrations in molasses-amended and -unamended reactors and in reactors switched to unamended conditions after establishing the active SCN^-^ degrading consortium. Five experiments were conducted over 790 days and genome-resolved metagenomics was used to resolve community composition at the strain level. A single *Thiobacillus* strain proliferated in all reactors at high loadings. Despite the presence of many *Rhizobiales* strains, a single *Afipia* variant dominated the molasses-free reactor at moderately high loadings. This strain is predicted to breakdown SCN^-^ using a novel thiocyanate dehydrogenase, oxidize resulting reduced sulfur, degrade product cyanate (OCN^−^) to ammonia and CO_2_ via cyanase, and fix CO_2_ via the Calvin-Benson-Bassham cycle. Removal of molasses from input feed solutions reproducibly led to dominance of this strain. Neither this *Afipia* strain nor the thiobacilli have the capacity to produce cobalamin, a function detected in low abundance community members. Although sustained by autotrophy, reactors without molasses did not stably degrade SCN^-^ at high loading rates, perhaps due to loss of biofilm-associated niche diversity. Overall, convergence in environmental conditions led to convergence in the strain composition, although reactor history also impacted the trajectory of community compositional change.

## Introduction

Thiocyanate (SCN^-^) is an unwanted toxic by-product found in a range of wastewater streams, including those from gold mines and coal coke processing [1]. The volume of freshwater contaminated by industrial processes is huge [2]. The requirement for treatment strategies is pressing, especially in countries such as South Africa and Australia, where gold recovery is an important economic activity and fresh water supplies can be very limited. There has been some progress with remediation of SCN^-^-containing effluents prior to discharge using mixed microbial consortia, but the systems are essentially treated as black boxes [3,4].

Some field reactors are inoculated with active laboratory-maintained consortia and continuously cultured with input of organic carbon, often in the form of molasses, and phosphate [3,5]. Previously, we characterized microbial communities derived from laboratory-maintained Activated Sludge Tailings Effluent Remediation (ASTER™, Outotec, South Africa) systems using a shotgun sequencing-based method that enabled recovery of genomes from many consortia members [6–8]. Different strain combinations proliferated under distinct operating conditions in these molasses-amended reactors, and the communities were very complex, with a reservoir of over 150 genotypes. Thiobacilli were the dominant community members at high SCN^-^ loadings and were inferred to be critical to SCN^-^ breakdown. Notably, the abundant thiobacilli were predicted to be autotrophs, bringing into question the need for added organic carbon compounds. Subsequently, [9] showed that distinct consortia could remove SCN^-^ from mining-contaminated solutions in the absence of added organic substrate and attributed SCN^-^ breakdown to three thiobacilli closely related to *T. thioparus* strain THI 111. In another recent study, laboratory bioreactors were inoculated from mine tailings and the autotrophically supported consortia found to breakdown SCN^-^ [10]. The currently known mechanisms for thiocyanate degradation are reviewed by Watts and Moreau [11].

Here, we describe a long-term study in which five reactors were inoculated from the same reactor as used by [6] and experiments conducted to test the hypothesis that the community could degrade SCN^-^ without the addition of organic carbon, specifically molasses, and that autotrophic thiobacilli would dominate the reactors at high SCN^-^ loadings. The reactors were driven to the same conditions via different routes to test whether historical contingency modulates the control of environmental conditions on community composition. We applied time-series genome-resolved metagenomics methods to investigate strain overlap (at the genotype level) among reactors that experienced a range of SCN^-^ loadings with or without molasses. Switching reactors from heterotrophic to autotrophic conditions consistently selected for the same SCN^-^ degrading strain from a background of substantial strain diversity. Insights from the genome of this strain include its pathways for degradation of SCN^-^ and breakdown products. Extensive genomic resolution of all communities enabled us to establish community convergence controlled by convergence of inputs and elucidated roles performed by other community members in this laboratory model system.

## Results and Discussion

### Experimental design and reactor chemistry

Five reactors (R1-R5) were inoculated from the same mixed microbial community and were operated continuously for 790 days. Residence times of 24 hours were maintained early in the experiments but were later lowered to 12 hours (**Table 1**). Each reactor experienced a different combination of SCN^-^ loading and molasses amendment, with consistent phosphate amendment over the full time course. The experimental design is shown in **Table 1** and **Figure S1** (also see Methods). As the experiments progressed, biofilm developed on all reactor internal surfaces. The biofilm was thick and dark brown in reactors receiving molasses and thin and creamy-white in reactors without molasses.

**Table 1.** Summary of experimental conditions. The table indicates continuously supplied thiocyanate feed concentrations, hydraulic retention times (HRT) and loading rates in molasses (M) or water-based (W) reactor systems (R1 - R5). The arrows in the Table indicate the points from which the experimental conditions in the reactors were held at a SCN^-^ feed concentration of 250 mg/L (0.36 mmol/L/hr).

| SCN feed concentration (HRT) | SCN loading rate (mmol/L/hr) | R1 | R2 | R3 | R4 | R5 |
| --- | --- | --- | --- | --- | --- | --- |
| 50 mg/L (24 hr) | 0.036 | M | M | M | M | W |
| 250 mg/L (12 hr) | 0.36 | M | M | M | M | W |
| 250 mg/L (12 hr) | 0.36 | M | ↓ | W | W | W |
| 500 mg/L (12 hr) | 0.72 | M |  | ↓ | W | W |
| 750 mg/L (12 hr) | 1.08 | M |  |  | W | W |
| 750 mg/L (12 hr) | 1.08 | M |  |  | W | W |
| 1500 mg/L (12 hr) | 2.15 | M |  |  | W | W |
| 2000 mg/L (12 hr) | 2.87 | M |  |  | W | W |

The SCN^-^ degradation rate consistently matched the SCN^-^ loading rate over the experimental period in three of the five reactors, demonstrated by negligible residual SCN^-^: R1 (which was subjected to an increasing SCN^-^ input concentration in the presence of molasses via the continuous feed, up to a maximum input SCN^-^ concentration of 2000 ppm), R2 (which was held at an input concentration of 250 ppm SCN^-^ with molasses), and R3 (which was held at an input concentration of 250 ppm SCN^-^ before and at all time points after molasses was removed from the input solution) (**Figure 1**). Thus, SCN^-^ accumulation did not challenge the survival of community members unable to degrade it. The sulfate concentrations in R1, R2 and R3 closely matched those predicted based upon reaction stoichiometry, given complete oxidation of all sulfide derived from SCN^-^ (**Table S1**). The concentrations of nitrogen compounds in the form of ammonia plus nitrate in R1, R2 and R3 also closely matched those predicted based on reaction stoichiometry, except in R1 after the SCN^-^ input concentration reached 1500 ppm. Notably, molasses-amended consortia outperformed consortia studied previously, which lost effectiveness at lower concentrations [7].

**Figure 1:**
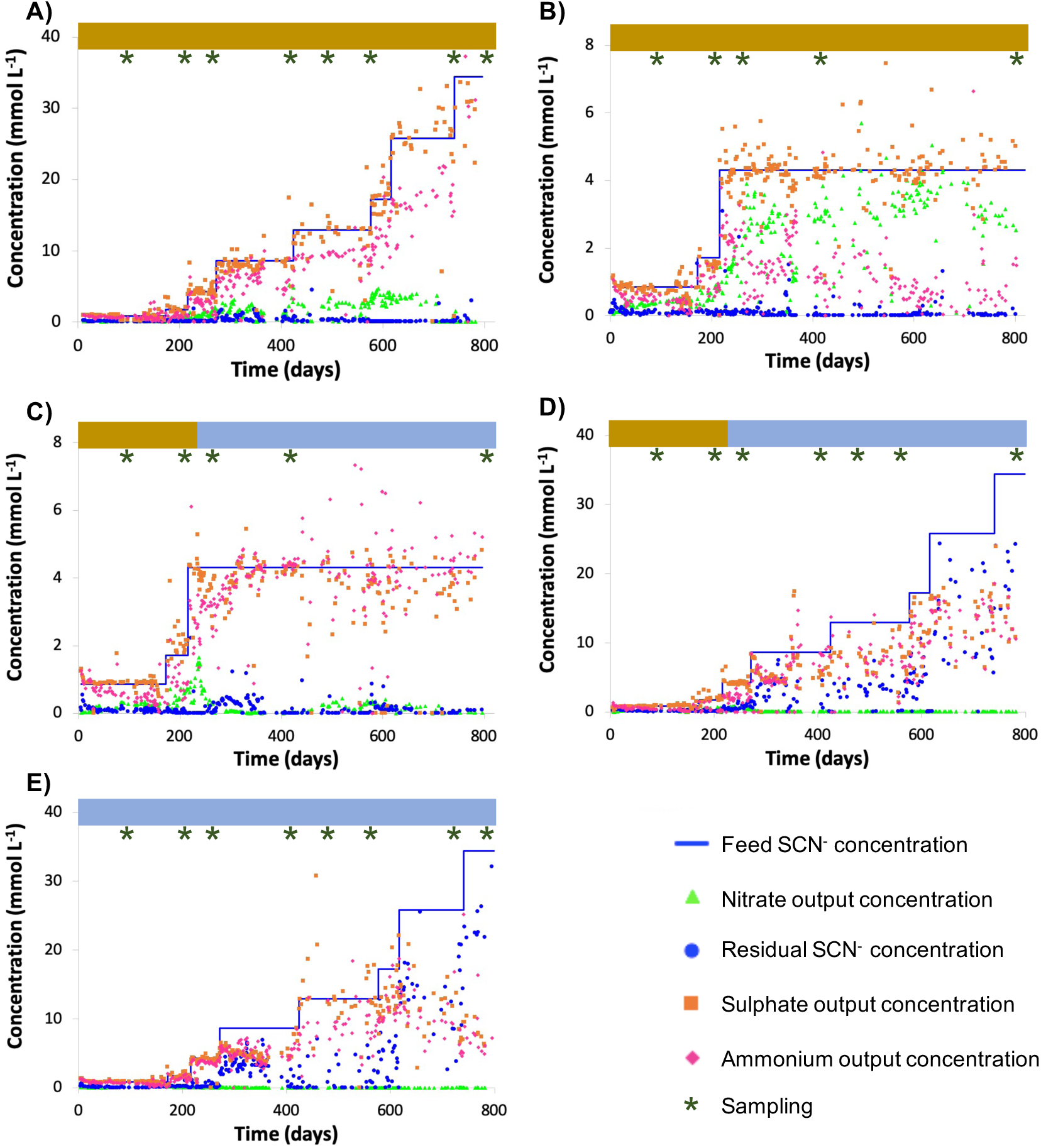
Chemical parameters for reactor 1 (R1; A), reactor 2 (R2; B), reactor 3 (R3; C), reactor 4 (R4; D) and reactor 5 (R5; E). Sampling time points for total DNA extractions are indicated above plots (brown: molasses, blue: no molasses).

R4 performed similarly to R1, R2 and R3 while maintained on molasses up to a SCN^-^ feed concentration of 250 ppm. After this point, molasses was removed from the input solution and the input SCN^-^ concentration was ramped from 250 to 2000 ppm. The SCN^-^ degradation rate declined and became erratic during the ramp. The toxicity of undegraded SCN^-^ in reactors at higher loadings could have impacted community composition. Although the sulfate and total nitrogen compound concentrations in R4 closely matched those predicted based upon reaction stoichiometry for complete breakdown of SCN^-^ early in the experiment, nitrate accumulation decreased at higher SCN^-^ input concentrations. This finding suggests a decline in abundance or activity of ammonia-oxidizing bacteria following accumulation of residual SCN^-^ in solution.

In the no-molasses reactor, R5, SCN^-^ was degraded completely up until the input concentration reached 250 ppm, consistent with findings from R3 and R4. Subsequently, the SCN^-^ degradation rate did not increase in direct proportion with increased loading rate, leading to incomplete SCN^-^ degradation. This was particularly true at high loading rates. Further, residual SCN^-^ concentrations fluctuated. While the ammonia production matched that predicted given the observed extent of SCN^-^ degradation, little or no nitrate was produced. Again, this suggests a decline in abundance or activity of ammonia-oxidizing bacteria at elevated SCN^-^ concentrations in the absence of organic feed.

### Genomic description of reactor communities

Genome binning of metagenomic data from 55 samples taken from the five reactors (see Methods) generated 1509 draft genomes, of which 956 were classified as near-complete (completion ≥ 90% and ≤ 5% contamination/redundancy) based on the predicted inventory of single copy genes from CheckM [12]. Given that metagenomic data were derived from samples collected from a series of related reactors at different time points, the genome dataset was dereplicated (see Methods), generating 232 genomes representative of dereplicated genome clusters, all of which correspond to bacteria. Of the dereplicated genomes, 168 were classified as near-complete. Between ∼70% and ∼80% of reads from each of the 55 metagenomic samples could be mapped back to the 232 genomes with high confidence (Methods and **Table S2**). Further, based on inventories of assembled single copy genes assigned to genomes, the abundant bacteria in all assemblies were represented by bins. Thus, we conclude that unmapped reads derived from low abundance organisms (some of which may be strain variants of more abundant organisms). The abundances of each of the 232 bacteria across the 55 samples are provided in **Table S3**.

We identified the abundant bacteria in R1 (ramped molasses reactor) and R5 (ramped water reactor) and calculated the abundances of these bacteria in all reactors (**Table 2A** and **Table S4**). We then predicted their relevant metabolic capacities (**Table 2B**). Notably, despite a wide variety of Rhizobiales strains, one specific genotype was highly abundant or dominated reactors maintained without molasses input, especially at intermediate to high SCN^-^ input concentrations (**Table 2** and **Figure 2**). This bacterium was not prominent in any molasses reactor from the current or prior studies, although it is closely related to the previously detected strain referred to as SCNPILOT_CONT_500_BF_Rhizobiales_64_17 [7]. Based on analysis of the 16S rRNA gene, this bacterium is related to organisms previously classified as either *Afipia* or *Bradyrhizobium*. We abbreviated the genome name as *Afipia*_64_1782. Given its importance in the community, we manually curated the genome representative of the dereplicated cluster from sample SCN18_10_11_15_R3_B, correcting many local assembly errors and filling scaffolding gaps using read sequences, extending and joining contigs (Methods). The essentially complete 3.549 Mbp genome has all expected single copy genes present in single copy and is in just 8 fragments (most fragments are terminated in unresolvable repeats).

**Table 2.**
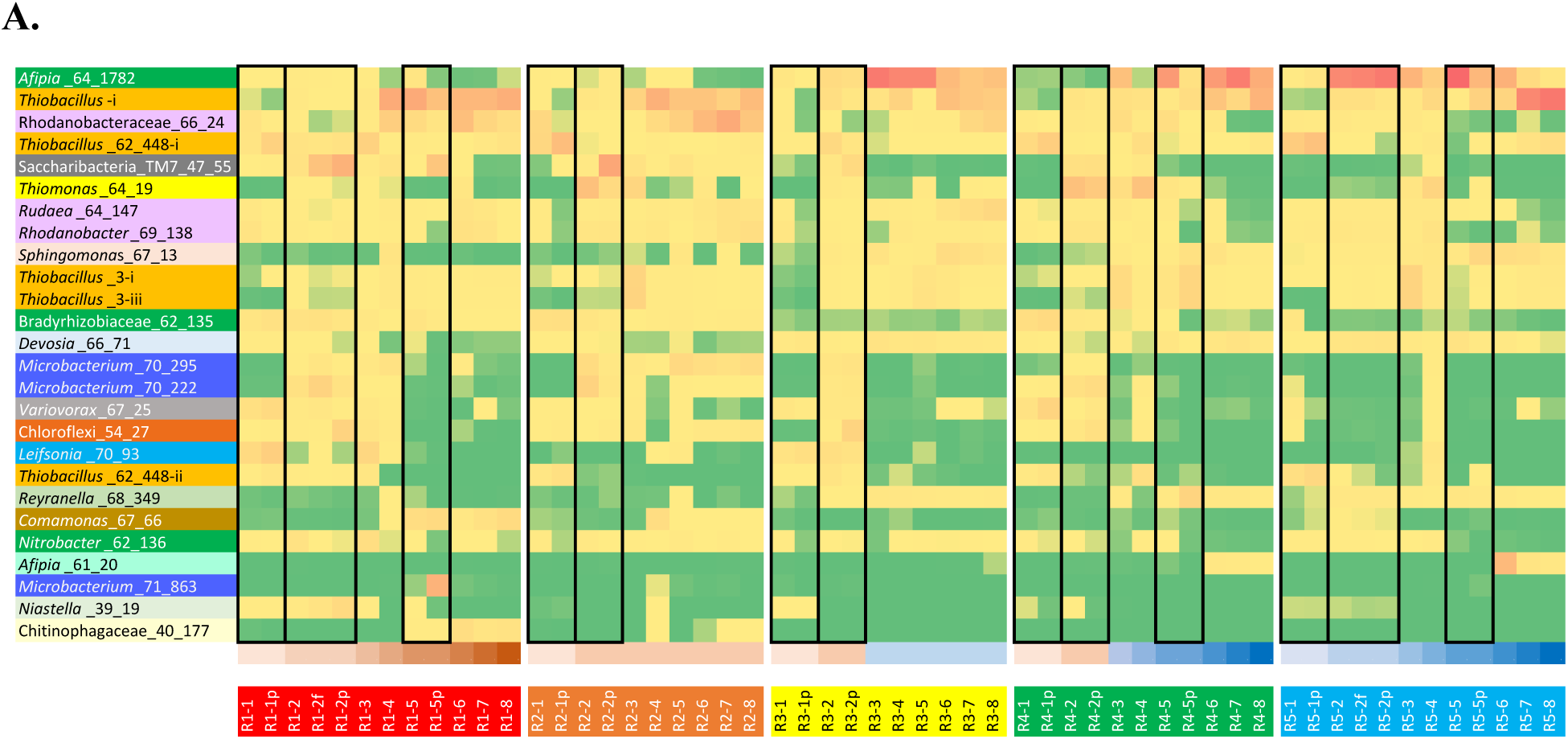
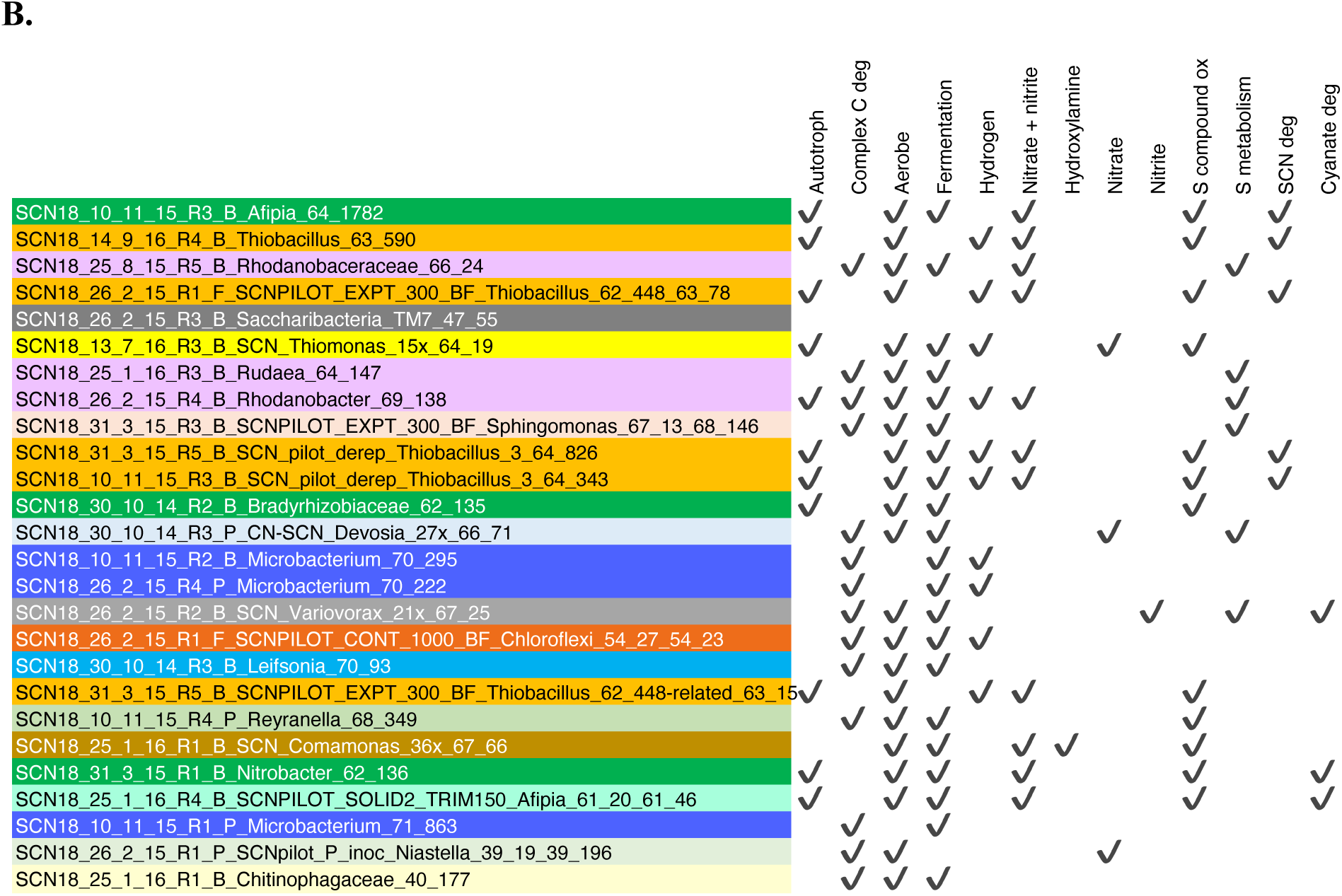
**A.** Pattern of abundance (high red, low yellow to green; see **Table S4**) of the 26 most abundant organisms (rows) in R1 (molasses SCN ramp reactor, shown in shades of brown at the bottom of the table) and/or R5 (SCN ramp reactor without molasses i.e. ‘water’ reactor, shown in shades of blue at the bottom of the table) across all reactors (names listed in colored boxes below). For the full names of bacteria see **B**. Black boxes group together samples from the same time point (biofilm and p=planktonic and f=floc sample). **B.** Overview of functionalities encoded in the 26 genotypes representing the most abundant organisms in R1 and/or R5.

**Figure 2:**
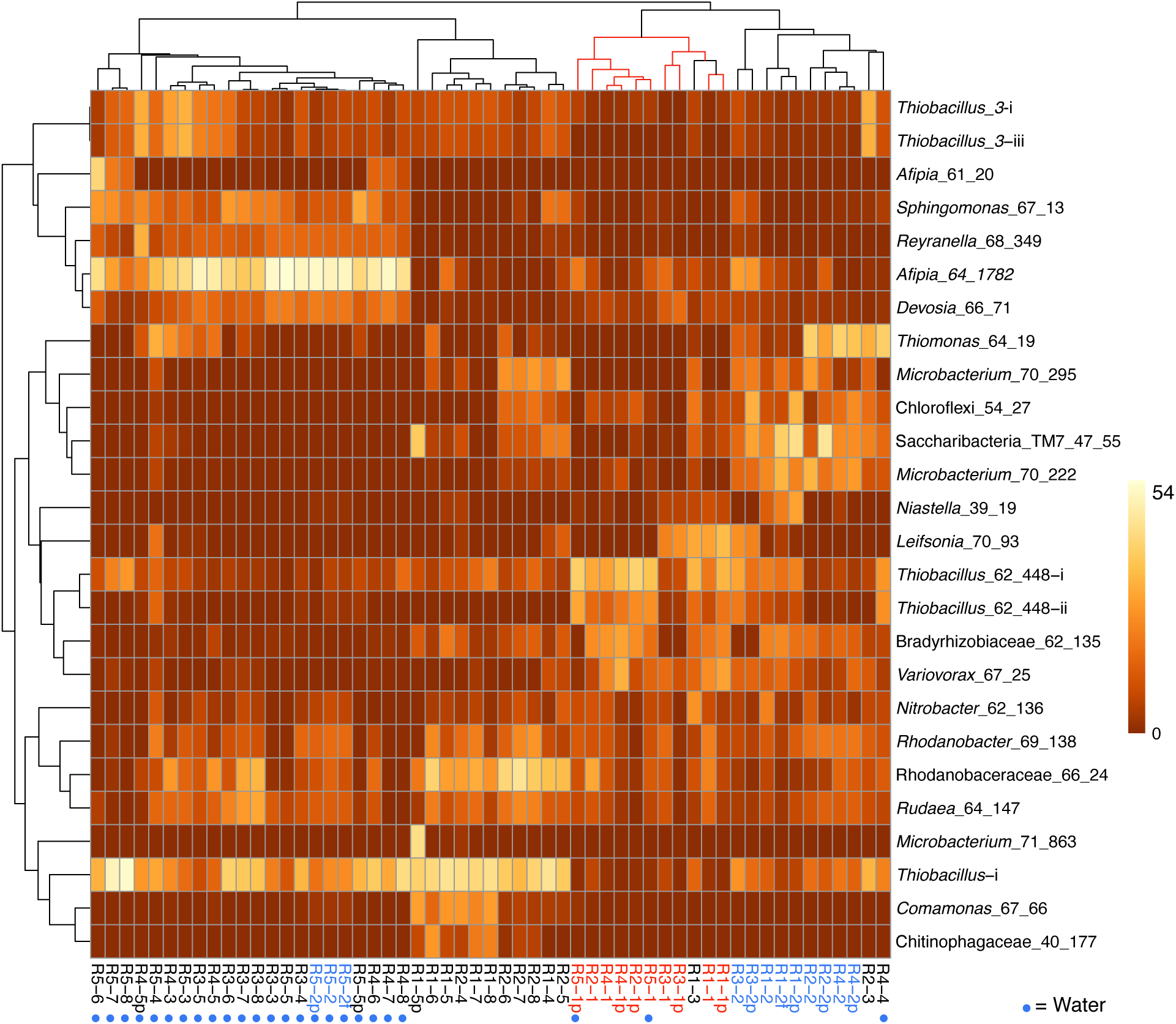
Heat map showing the clustering of the patterns of abundance of the 26 most abundant organisms in reactor 1 (molasses SCN ramp reactor) and/or reactor 5 (water SCN ramp reactor). The plot is log(x+1) transformed with no row centering and includes some planktonic (p) and floc (f) samples as well as biofilm samples. Red text and lines indicate the first samples from the five reactors and blue text indicates the second samples from these reactors. See **Table 2A** for the pattern of abundance of these organisms over samples and reactors and **Table 2B** for the full names of all genotypes. Note that, with the exception of the starting communities (Rx-1) and R4-4, communities in the reactors not receiving molasses cluster together.

Based on the prevalence of abundant bacteria in the five reactors, the molasses-free reactor communities cluster together and away from communities from reactors with molasses in the input solution (**Figure 2**). Thiobacillus_3-i and Thiobacillus_3-iii, which are closely related to strains abundant previously in molasses-amended reactors [7], were almost undetectable in initial samples but they proliferated in later samples, especially those from reactors not receiving molasses. *Thiobacillus*_63_590, shown as *Thiobacillus*-i in **Figure 2** and **Table 2A**, was barely detectable at the first time point but generally increases in abundance with increasing SCN^-^ input concentration and is the dominant organism in R1 and R2 at later time points.

### Predicted metabolism and capacity for SCN^-^ degradation

Given the proliferation of *Afipia*_64_1782 in reactors without molasses that achieved SCN^-^ degradation, we searched the genome for genes capable of degrading SCN^-^ and its breakdown products. We identified a 489 aa protein (SCN18_10_11_15_R3_B_scaffold_20_388) that was initially annotated as thiocyanate hydrolase, but the three subunits expected (*scnABC* of the carbonyl sulfide pathway) were not present. Further analyses (Methods) indicated that the gene encodes thiocyanate dehydrogenase of the cyanate pathway [13] (**Figure 2**). This identification was strongly supported by selection of threading templates using I-tasser [14] as well as by SWISS-MODEL [15] (33.7% identity, top match to PDB 5f75, with four Cu ligand sites identified although the GMQE score is below expectation relative to the value expected for experimental structures of similar size) and Phyre^2 [16]^ (template fold library ID c5f30B, thiocyanate dehydrogenase from *Thioalkalivibrio paradoxus*). The sequence is closely related to that of mesophilic alphaproteobacteria strain THI201, which was shown to degrade thiocyanate and is also well modeled by the PDB 5f75 structure. The genome also encodes a cyanate lyase/cyanase (SCN18_10_11_15_R3_B_scaffold_20_38), which could convert the cyanate (OCN^-^) produced by thiocyanate dehydrogenase to ammonia and CO_2_. Also identified is carbonic anhydrase, so the bacterium may use the cyanase to convert cyanate to carbamate (for use in pyrimidine metabolism). The *Afipia*_64_1782 also has sulfide dehydrogenase, sulfide:quinone oxidoreductase and genes of the SOX pathway to oxidize sulfide released by SCN^-^ degradation (as well as genes for sulfide assimilation). Although apparently unable to oxidize ammonia (produced by thiocyanate dehydrogenase and cyanase), the *Afipia*_64_1782 genome encodes a nitrate/nitrite transporter, the capability to convert nitrite to nitrate, reduce nitrate (NO-forming), as well as nitric oxide reductase and nitrous oxide reductase (thus, it is predicted to be capable of complete denitrification). Given that essentially all N predicted to be produced from SCN^-^ degradation is accounted for by ammonia in reactors without molasses (**Figure 1**), this bacterium may not be reducing nitrate or nitrite *in situ*.

The abundant bacteria in R5 are presumably autotrophs, given the absence of added organic carbon. *Afipia*_64_1782 encodes two genes annotated as RuBisCO (confirmed to be the small and large subunits of Form ID, based on phylogenetic analysis of the large subunit) and other genes of the Calvin-Benson-Bassham (CBB) cycle for CO_2_ fixation. The *Afipia*_64_1782 is predicted to be capable of aerobic growth. Its genome encodes a complete glycolysis/gluconeogenesis pathway, TCA cycle as well as the glyoxylate cycle, genes of the pentose phosphate pathway, including those required to synthesize precursors for nucleic acid synthesis and for purine and pyrimidine biosynthesis. The genome also encodes genes for production of trehalose, starch/glycogen and aminosugars including UDP-N-acetylmuramate precursor for peptidoglycan production and for peptidoglycan production. Also identified are genes for synthesis of a Gram negative cell envelope, acetate production via acetyl-P and interconversion of pyruvate and lactate. Interestingly, although the genome appears to encode genes for the biosynthesis of folate, NAD, pantothenate, CoA, thiamine, riboflavin, and probably biotin, this bacterium appears to be unable to produce cobalamin.

As expected based on prior work [6–8,10], various thiobacilli were prominent in the reactors. However, *Thiobacillus*_63_590, which was highly abundant especially in the molasses reactors and in the molasses-free reactor at the highest SCN^-^ input concentrations (*Thiobacillus*-i in **Table 2A**) is distinct from strains/species abundant in prior studies. Similar to previously described abundant thiobacilli in SCN^-^ degrading reactors, this bacterium is an autotroph with both Forms I and II RuBisCO. Its genome encodes the three subunit thiocyanate hydrolase, the necessary transporters and cyanase, carbonic anhydrase to form bicarbonate for assimilation of cyanate into carbamate. It also has genes that may breakdown organic sulfur compounds, genes of the SOX and the reverse DSR pathways for inorganic sulfur compound oxidation, for uptake of nitrate and nitrite, complete denitrification and recovery of ammonia from nitriles. This bacterium has genes for biosynthesis of nucleic acids, a complete pathway for synthesis of peptidoglycan, isoprenoid precursors via the MEP pathway and the ability to store carbon compounds as starch. Like *Afipia*_64_1782, it cannot synthesize cobalamin *de novo* but it is predicted to synthesize other cofactors, including thiamine, biotin and riboflavin.

Overall, the reactors contain numerous other bacteria that likely confer community-relevant functions. Several other bacteria abundant in the molasses-free reactor that proliferated at moderate to high SCN^-^ input concentration also have Form I and / or Form II RuBisCOs and are predicted to fix CO_2_ via the CBB cycle (**Table 2B**). Three relatively abundant bacteria belonging to the Rhodanobacteraceae (pink names in **Table 2B**), *Sphingomonas* and Chloroflexi are predicted to have genomes that encode numerous enzymes involved in degradation of complex carbon compounds. Many genomes of coexisting *Mesorhizobium* and *Bosea, Reyranella*, various other Alphaproteobacteria (especially Rhizobiales), as well as *Pseudonocardia*, organisms that are present at relatively low abundance in the communities, appear to have genes required to produce cobalamin. Only two bacteria, Rhizobiales_68_13 and Verrucomicrobia_61_8_61_17, are predicted to have the capacity to fix N_2_. However, many bacteria are predicted to have various combinations of genes for nitrate, nitrite, nitric oxide and nitrous oxide reduction. Similarly, many have genes for sulfide/sulfur compound oxidation via SOX genes, but the thiobacilli appear to be the main group using the reverse dissimilatory sulfite reductase pathway. Only a few bacteria, including some Actinobacteria (various *Microbacterium* species) and Saccharibacteria (TM7) are predicted anaerobes. Similar abundance patterns of the TM7 and *Microbacterium ginsengisoli* in R2 and R3 may indicate a Saccharibaceria (TM7)-actinobacterial host association similar to that reported previously for oral Saccharibacteria [17]. The clustering in **Figure 2** supports this inference, although the pattern is not apparent in all analyses (**Figure S2**).

As expected based on the chemical data that show complete oxidation of sulfide derived from thiocyanate degradation to sulfate, many of the more abundant non-SCN^-^ degrading bacteria in all reactors have genes for sulfur compound oxidation (**Table 2B**). However, organisms potentially responsible for conversion of ammonia to nitrate that were detected primarily in molasses amended reactors are less apparent. In fact, *amoA,B,C* genes involved in ammonia oxidation were not identified in any of the abundant bacteria (**Table 2B**). Similarly, nitrifying microbes were not identified in the study system of Watts et al. [10], despite the potential value of ammonium as an energy source. In the study of Kantor et al. [7], it was inferred that *Nitrosospira* were responsible for ammonia oxidation in molasses-amended reactors, based on unbinned sequences. Here, we identified three draft genomes of relatively low relative abundance *Nitrosospira* strains with genes annotated as *amoA,B,C*. One genome for a strain related to *Nitrosospira multiformis* also encodes hydroxylamine oxidoreductase (HaoA,B), as occurs in *Nitrosomonas europaea [18]*. Interestingly, *Comamonas* and Burkholderiales genomes also have HaoA,B. All HaoA predicted proteins have the anticipated 8 CxxCH motifs. Genes for ammonia oxidation were not identified in the *Comamonas* and Burkholderiales genomes. *Comamonas* were only relatively abundant at later time points in R1 and R2, but this may be unrelated to the presence of HaoA,B.

The strains of *Nitrosospira* implicated in ammonia oxidation switch in relative abundance over time. The strain represented by SCN18_31_3_15_R1_B_Nitrosospira_56_42 is most prominent at the start of the experiments as well as at low to intermediate SCN^-^ input concentrations, regardless of whether molasses was added. However, *N. multiformis*, which is at very low abundance in all reactors at the first time point, increases in abundance in molasses-amended reactors at longer times, especially at higher SCN^-^ loadings. The proliferation of this strain only in molasses-amended reactors may explain the very low production of nitrate in reactors without added molasses. Overall, the findings suggest that the activity of a variety of *Nitrospira* strains, some apparently at quite low abundance, are responsible for nitrification when it occurred in the reactors. Some ammonia released by SCN^-^ degradation is presumably assimilated for biomass production and, as noted above, a fraction of SCN^-^-derived N partitioned into cyanate may also be assimilated.

### Community shifts with reactor conditions and time

We used principal component analysis (PCA) to investigate whole community compositions over time and as a function of reactor conditions. At the first time point, when R1-R4 had received molasses and R5 had been maintained over the startup period of 117 days without molasses at a low SCN^-^ feed concentration of 50 ppm and a high residence time of 48 - 24 h, the communities generally clustered together. However, the planktonic fraction of R5 was displaced towards that of the R5 consortia sampled at time point 2 after ramping the input solution from 50 ppm to 250 ppm and reducing the residence time to 12 h (**Figure S3**). This dramatic shift^-^ occurred within an additional 119 days. Differences in community composition in R1-R4 biofilm samples at time point 2 (despite experiencing the same conditions) may be due to stochastic processes during community development or may reflect within-reactor heterogeneity. However, within-reactor heterogeneity is unlikely to account for differences in the planktonic communities, as the reactors were well mixed (**Figure S3**).

At both the first and second time points, the biofilm, planktonic and samples of flocculated biomass (“flocs”) from each reactor are typically more similar to each other than to communities in other reactors (and planktonic and floc samples are essentially indistinguishable). PCA plots for individual reactors indicate some separation of biofilm and planktonic samples (**Figure S4**). Of the relatively abundant bacteria in R1, strong partitioning into the planktonic relative to biofilm fraction occurred only for *Microbacterium*_71_863 and Saccharibacterium at time point 5. This Saccharibacterium is also enriched in the planktonic fraction in R2 (second time point). Interestingly, *Afipia*_64_1782, which dominates molasses-free reactors and is at relatively low abundance in R1, is partitioned into the biofilm fraction of R1. R4 at time point 5 shows strong enrichment of a different microbacterium (*Microbacterium_71_138*) in the biofilm compared to planktonic fraction, consistent with the prediction that it is an anaerobe (**Table 2B**). The only strong fractionation in the molasses-free reactor, R5, occurred at time point 5 and involved partitioning of three abundant thiobacilli and a moderately abundant WPS-2 (candidate phylum Eremobacteraeota [19]) into the planktonic fraction.

Independent PCA analyses of R1, R4 and R5 (**Figure S4**) show step-like community compositional change as reactor SCN^-^ input concentrations were ramped. In contrast, the later timepoints in both R2 and R3 (timepoints 4 to 8 at constant input SCN^-^ concentration of 250 ppm) cluster together. Both of these observations would be expected if input SCN^-^ concentration is the primary compositional control within each reactor. Given these results, we compared the community compositions of R2 and R3 (**Figure 4**) and of R1, R4 and R5 (**Figure 5**). The PCA plot for R2 and R3 illustrates dramatic divergence attributable to termination of molasses addition. Notably, this separation occurred within 33 days of removal of molasses from the input solution for R3 (**Figure 4**). In detail, communities at timepoints 3, 4 and 5 separate from those at timepoints 6, 7 and 8 in both R2 and R3, a result that suggests ongoing evolution in community composition independent of SCN^-^ input concentration. The PCA plot for R1, R4 and R5 (**Figure 5A**) also clearly shows differences in community composition in molasses-free (R5) compared to molasses-amended reactors after reactors were ramped from SCN^-^ input concentrations of 50 to 250 ppm. The shift in R5 is generally tracked by communities in R4 after molasses was removed from input solution at time point 3. However, unexpectedly, R4-4 clusters with R4-2, possibly due to the unusually high abundance of *Thiomonas* in R4-4 (Tm, **Figure 5B**). Overall, the clustering of communities at moderate to high input SCN^-^ concentrations is driven by the proliferation of *Afipia*_64_1782 strain in the water reactors and one specific *Thiobacillus* strain in both molasses-free and molasses reactors (**Figure 5B**).

**Figure 3.**
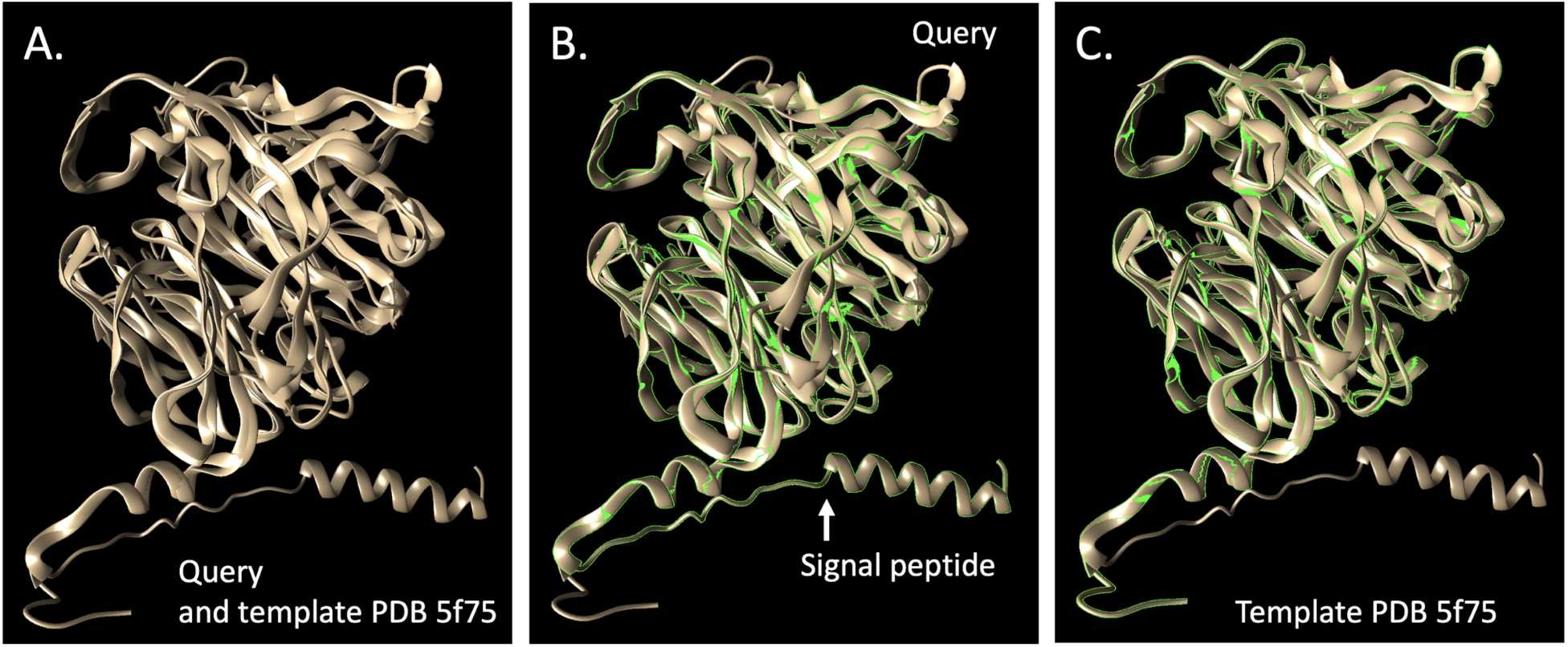
Comparison of the putative thiocyanate dehydrogenase from *Afipia*_64_1782 and the most closely related protein, Model 5f75, in the database (PDB) structure. **B.** shows the query and **C**. the template. The TM-score of the structural alignment between the query and 5f75 was 0.901, compared to the second choice score (PDB 2ece, a hypothetical selenium-binding protein from *Sulfolobus tokodaii*) of 0.608.

**Figure 4:**
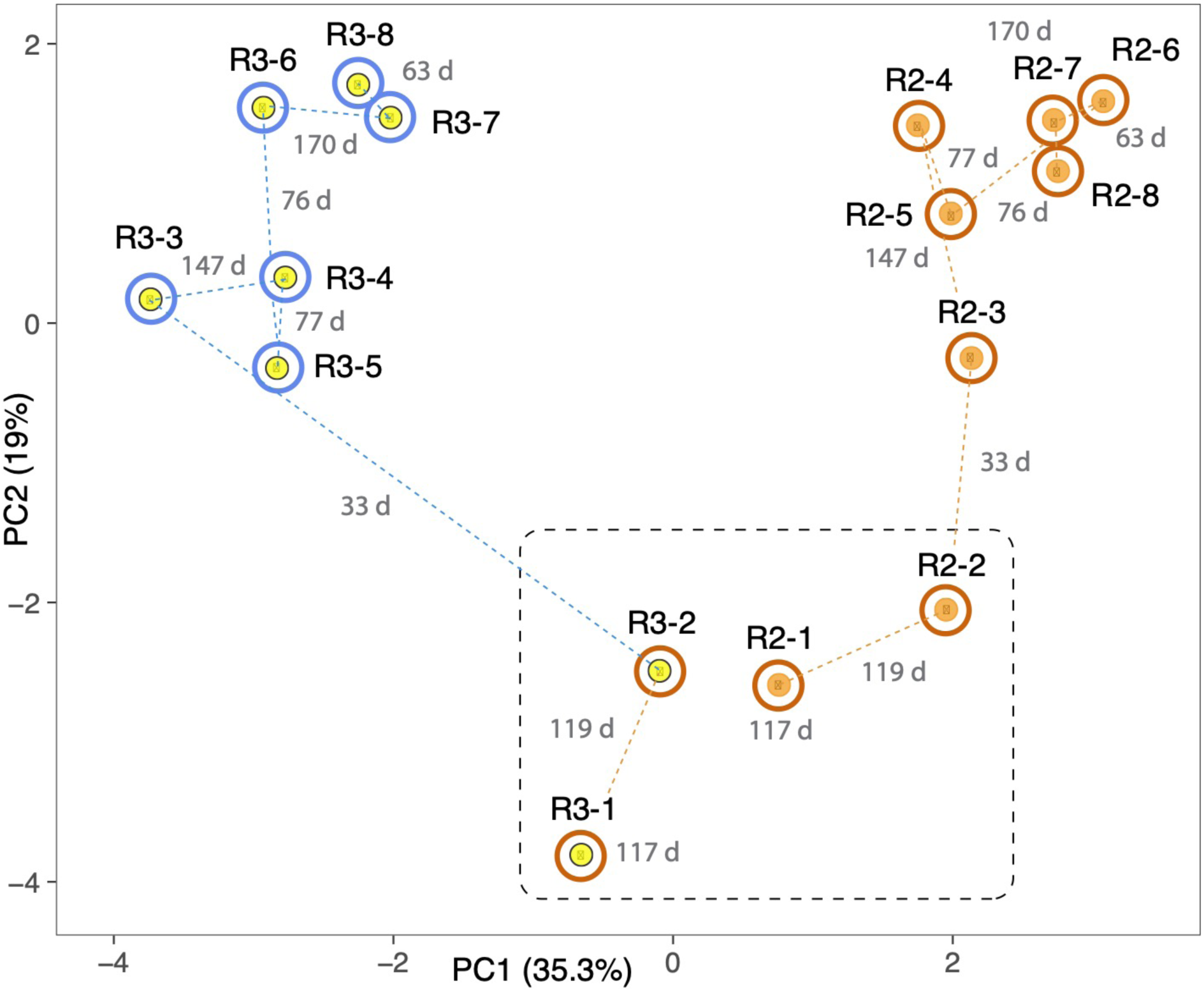
Principal component analysis based on the relative abundances of 232 organisms (accounting for the most abundant 70-80% of the community) in reactors 2 (orange dots) and 3 (yellow dots). Both reactors were ramped to 250 ppm SCN and held at this concentration for the duration of the experiment. Outer circles distinguish reactors receiving molasses (brown) from those receiving water (blue) at the time of sampling. The dashed black line separates samples taken prior to the switch of reactor 3 to water. Dashed lines indicate sequential samples and numbers in grey indicate the number of days (d) between sampling times. Note the strong and rapid deviation of community after the switch to water in reactor 3. Samples 6, 7 and 8 in both reactors group away from 3, 4 and 5, suggesting that the community shift was incomplete at the middle time points.

**Figure 5.**
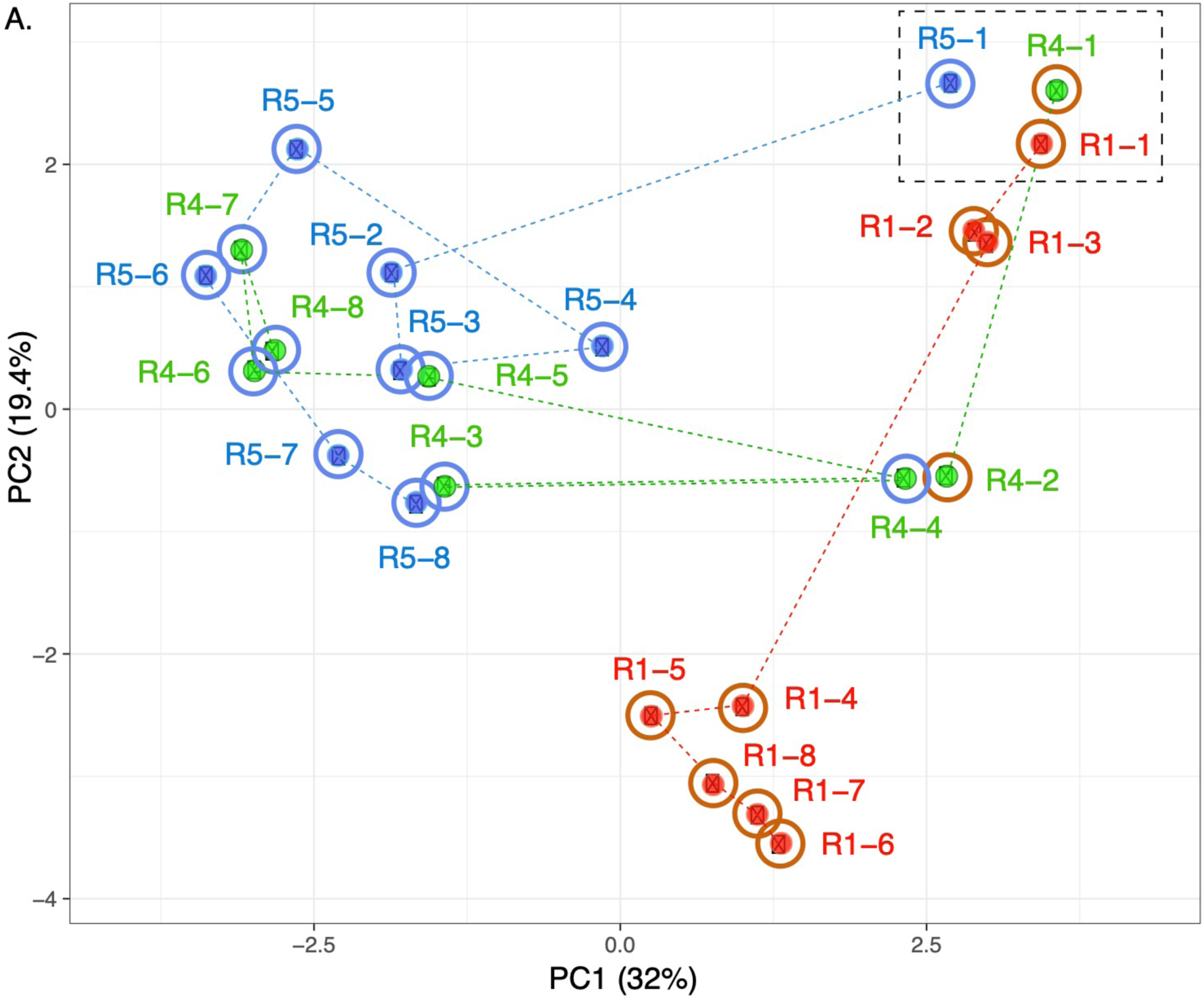
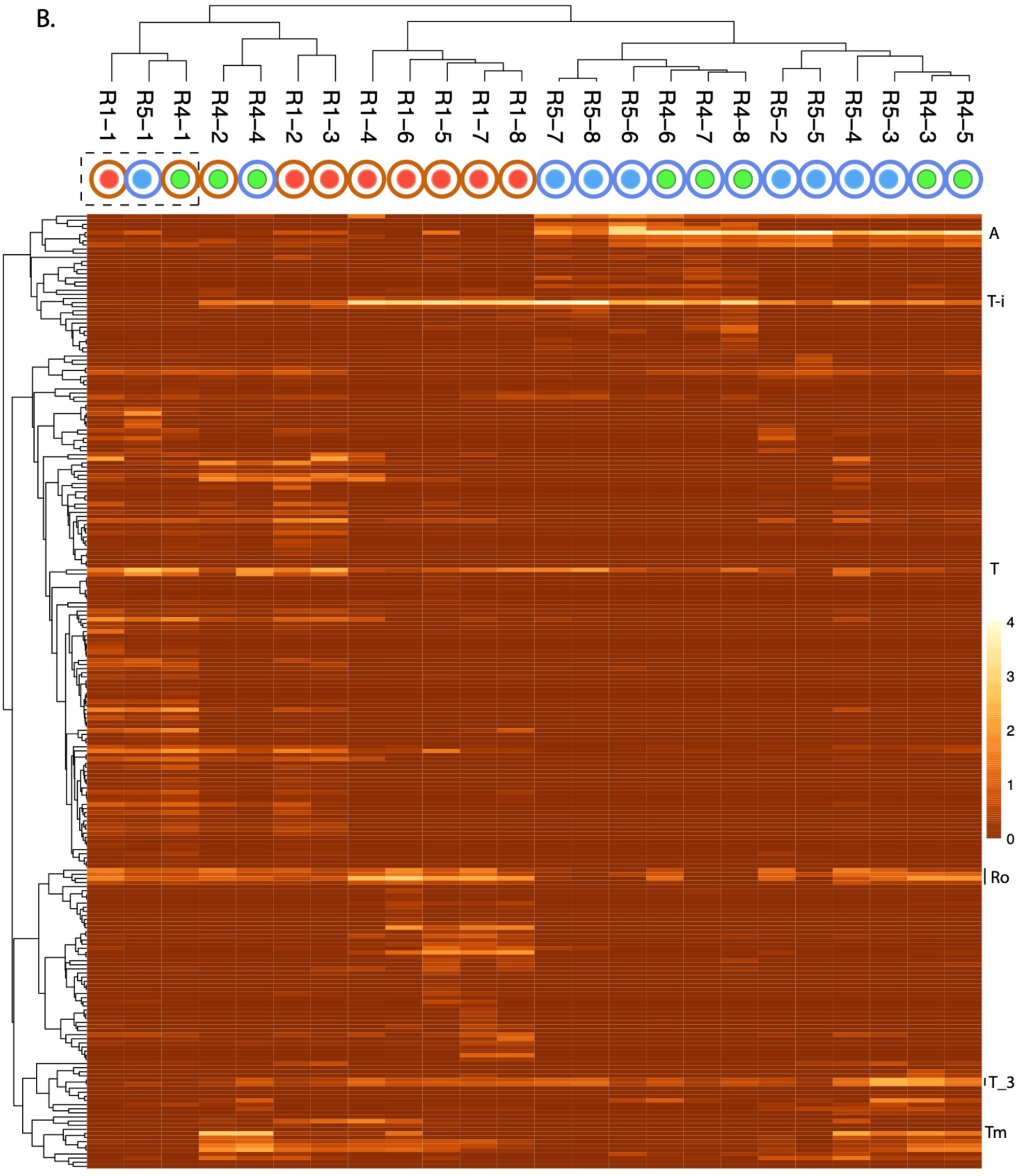
Principal component analysis based on the relative abundances of 232 organisms (accounting for the most abundant 70-80% of the community) in reactors 1 (red dots), 4 (green dots) and 5 (blue dots). The dashed box indicates the first samples from each reactor. Note the dramatic divergence of R5 after ramping from 50 to 250 ppm SCN^-^ in the absence of molasses, and the convergence of R4 with R5 after termination of molasses input. **B**. Heat map based showing clustering of samples based on the abundance of each of the 232 organisms. The proliferation of *Afipia*_64_1782 (A) separates most molasses-free (blue outer circle) from molasses-amended (brown outer circle) communities. Other abundant organisms include *Thiobacillus*-i (T), two strains related to *Thiobacillus*_3 (T_3), several Rhodanobaceraceae (Ro). *Thiomonas* (Tm) that is unusually abundant in R4-4 and may contribute to its unexpected clustering with R4-2.

## Conclusions

By combining a substantial set of long term-reactor-based experiments with resolution of the strain composition achieved via genome-resolved metagenomics we demonstrate consistent (across reactors) and largely reproducible (across experiments) selection for specific bacteria capable of autotrophic growth and thiocyanate degradation from a background of considerable strain diversity. Nonetheless, there is evidence for ongoing evolution in community composition in reactors maintained under constant conditions and some divergence in reactors experiencing the same conditions may be evidence for historical contingency effects. The reactors clearly diverge based on the presence or absence of added organic substrate in the form of molasses. Interestingly, despite the association of key genes for SCN^-^ degradation with autotrophs in both reactors and the prominence of a variety of autotrophs in molasses-free reactors, SCN^-^ degradation is destabilized in the absence of molasses, especially at high input SCN^-^ concentrations. We attribute this to the importance of heterotrophs that provide community-essential functions, including production of cobalamin, and creation of biofilm and its associated niche diversity.

## Methods

### Reactor Setup, Inoculation, and Operation

Five 1 L water-jacketed continuous stirred tank reactors (CSTRs; Glass Chem, South Africa) were inoculated with homogenized biofilm and planktonic samples harvested from a long-running SCN^-^ −degrading stock reactor at the University of Cape Town. These reactors, R1 - R5, were stirred with a pitched-blade impeller at 270 rpm and sparged with filtered air at 900 mL/min. Four reactors (R1 - R4) were initially fed 150 mg/L molasses and 0.28 mM KH_2_PO_4_to provide supplemental nutrients, while the fifth reactor (R5) was only fed deionized water containing 0.28 mM KH_2_PO_4_. All the reactors were further supplemented with KSCN as detailed below. The respective reactor feeds also contained increasing amounts of KOH to modulate reactor pH as necessary and small amounts of 5 N KOH were added dropwise directly to reactors if observed pH was ≤ 6.5.

The reactors were run in batch-fed mode for 4 days, before all the reactors were switched to continuous feeding at a residence time of 48 h. The hydraulic retention time (HRT) of the reactors was sequentially lowered from 48 to 12 h, over a period of 131 days, and then maintained at 12 h, while the feed concentration of SCN^−^ in the feed was increased stepwise over the remainder of the experimental period. The reactors were allowed to stabilize between each increase in the loading of SCN^-^ to reach steady state. The molasses was dropped from the feed to reactors 3 and 4 (R3 and R4) at Day 239 for the duration of the experiment. The SCN^-^ feed concentration to R2 and R3 was maintained at 250 mg/L, from day 216, for the remainder of the experiment.

Samples of planktonic cells and biofilm attached to the walls of the bioreactor were initially sampled at a 24 h HRT and SCN^-^ feed concentration of 50 mg/L (SCN^-^ loading rate: 0.04 mmol/L/h). The remaining samples were taken at a 12 h HRT and SCN^-^ feed concentrations of 50 to 2000 mg/L (SCN^-^ loading rates of 0.36 to 2.87 mmol/L/h).

### Sampling

Samples of biomass from each reactor were taken for metagenomic sequencing just before increases in feed concentration. Approximately 0.5 g (wet-weight) of biofilm was scraped from the wall of each reactor with sterile spatula and stored at −60 °C. Paired samples of planktonic biomass were collected by filtering approximately 300 mL of the liquid phase from each reactor through sterile Whatman paper, to remove small flocs and clumps of biofilm, before the planktonic cells were recovered onto sterile 0.22 μm filters. Biomass was gently washed off the filter with sterile water, harvested by centrifugation (13000 rpm for 10 min at room temperature) and stored at −60 °C until further analysis. Filtered media was returned to the reactor to maintain chemical continuity.

### Analysis of Solution Chemistry

Bulk liquid was sampled daily for chemical analysis, filtered through a sterile 0.22 μm syringe filter, pH analyzed, and frozen at −20 °C until further analysis. Residual SCN^−^ concentration was measured using High Performance Liquid Chromatography (HPLC) as described previously. Ion chromatography (IC) was performed to quantify ammonia, nitrate and sulphate on a Dionex ICS1600 system (Thermo Scientific). Ammonia was quantified using a Dionex Ionpac CS12A column, while sulphate and nitrate were quantified using a Dionex Ionpac AS16 column.

### DNA Extraction and Sequencing

DNA was extracted from frozen biomass using a NucleoSpin soil genomic DNA extraction kit (Machery-Nagel, Germany) with the inclusion of a repeated extraction step, according to the manufacturer’s instructions.

### Metagenomic Assembly, Binning, and Annotation

The metagenomic data processing of paired 150 bp read datasets followed methods reported previously [7] up to the genome binning step. To generate genome bins, differential abundance information was calculated for the scaffolds of each sample using the abundance of scaffolds across all samples. The reads of each sample were mapped against the scaffolds of each sample using BBMap (https://sourceforge.net/projects/bbmap/) at 98% average nucleotide identity. The series coverage information was used to create bins using ABAWACA [20], CONCOCT [21], MaxBin2 [22], and MetaBAT2 [23]. Bins were also manually generated by leveraging information derived from matching against a dereplicated set of genomes from prior research SCN^-^-degrading communities [6–8] and UniProt [24]. The resulting taxonomic profile and other “manual binning” signals (GC content, coverage, and single copy gene inventory were integrated for human-guided binning. A set of optimal bins for each sample was created from the combined automated and manual bins using DASTool [25]. All of the DASTool bins were dereplicated using dRep to create a near-final draft genome set. The resulting dereplicated bins were further manually refined using the aforementioned “manual binning” signals. The manually refined bins were dereplicated with dRep to create a final dereplicated set of bins for all subsequent analyses. Relatives abundances for each genome were calculated by normalizing the coverage of each genome by the total coverage in each sample and then multiplying this fraction by the percentage of reads mapped in each sample.

The genome of *Afipia*_64_1782 was selected for manual curation [26]. This involved mapping of reads to the *de novo* assembled sequences, identification (based on regions of no perfect read support) and resolution of local mis-assemblies, gap filling, contig extension and joining based on perfect overlap of a region spanned by paired reads. Curation was terminated when sequences ended in repeats that could not be uniquely spanned using short Illumina read data.

Functional predictions for all genes in the final dereplicated reference genome set were established as previously described [7] but with additional constraints from an HMM-based analysis [27]. For specific genes of interest, analysis included prediction of signal peptides, domain architecture (e.g., by Interpro). For thiocyanate dehydrogenase, the best model in PDB was selected as the best choice by i-Tasser [14]. The comparison was visualized by UCSF Chimera [28].

### Principal coordinate analysis

Heatmaps and PCA plots were generated using ClustVis [29]. Original values are ln(x + 1)-transformed. No scaling is applied to rows; SVD with imputation is used to calculate the principal components. X and Y axes show principal component 1 (PC1) and principal component 2 (PC2) that explain the percentage (%) of the total variance, respectively.

## Acknowledgments

This research was supported by a grant from the National Science Foundation (USA) to JFB (EAR-1349278) and an NSF Graduate Fellowship to RK and DST/NRF of South Africa SARChI Chair in Bioprocess Engineering (UID 64778) to STLH and, DST/NRF Competitive Support for Unrated Researchers (CSUR) Grant (UID 111713) and Research Career Advancement Fellowship (UID 91465) to RJH. We thank Shufei Lei, Katherine Lane, Brian Thomas, and the QB3 Vincent J. Coates Genomics Sequencing Laboratory for research support.

## Author contributions

The five reactor experiments were designed by RH, STLH and RK, with input from JFB, and conducted at CeBER by RH and FK. Chemical analyses were performed by FK and RH, with input from STLH. RH extracted the DNA, which was sequenced at the UC Berkeley QB3 facility. RS conducted the metagenomic assembly, performed functional annotations, and performed bioinformatics analysis for automated binning using previous genomes from the study system. JFB performed manual genome binning, bin curation and genome curation. JFB, RS, STLH and RH analyzed the data and JFB and RH wrote the manuscript, with input from RS and RK. All authors commented on and approved the final version of the paper.

## Supplementary Information

### Supplementary Figures

**Figure S1.**
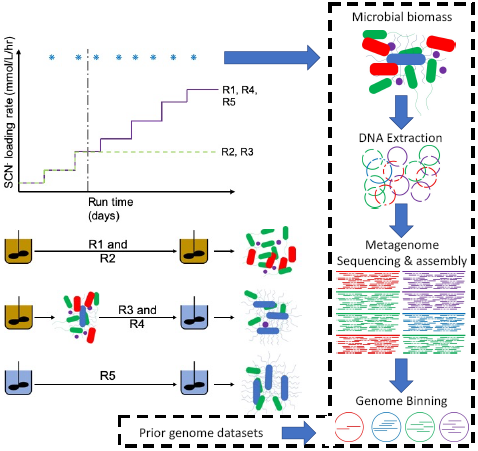
Diagram summarizing the operation and sampling of the five SCN^-^reactors over the course of the experimental period. A) Shows the experimental SCN-feed loading regime over time, while (B) shows the five SCN-degrading reactors in operation. The vertical dotted lines and asterisks show the points at which biomass was sampled for subsequent total genomic DNA extractions and sequencing.

**Figure S2.**
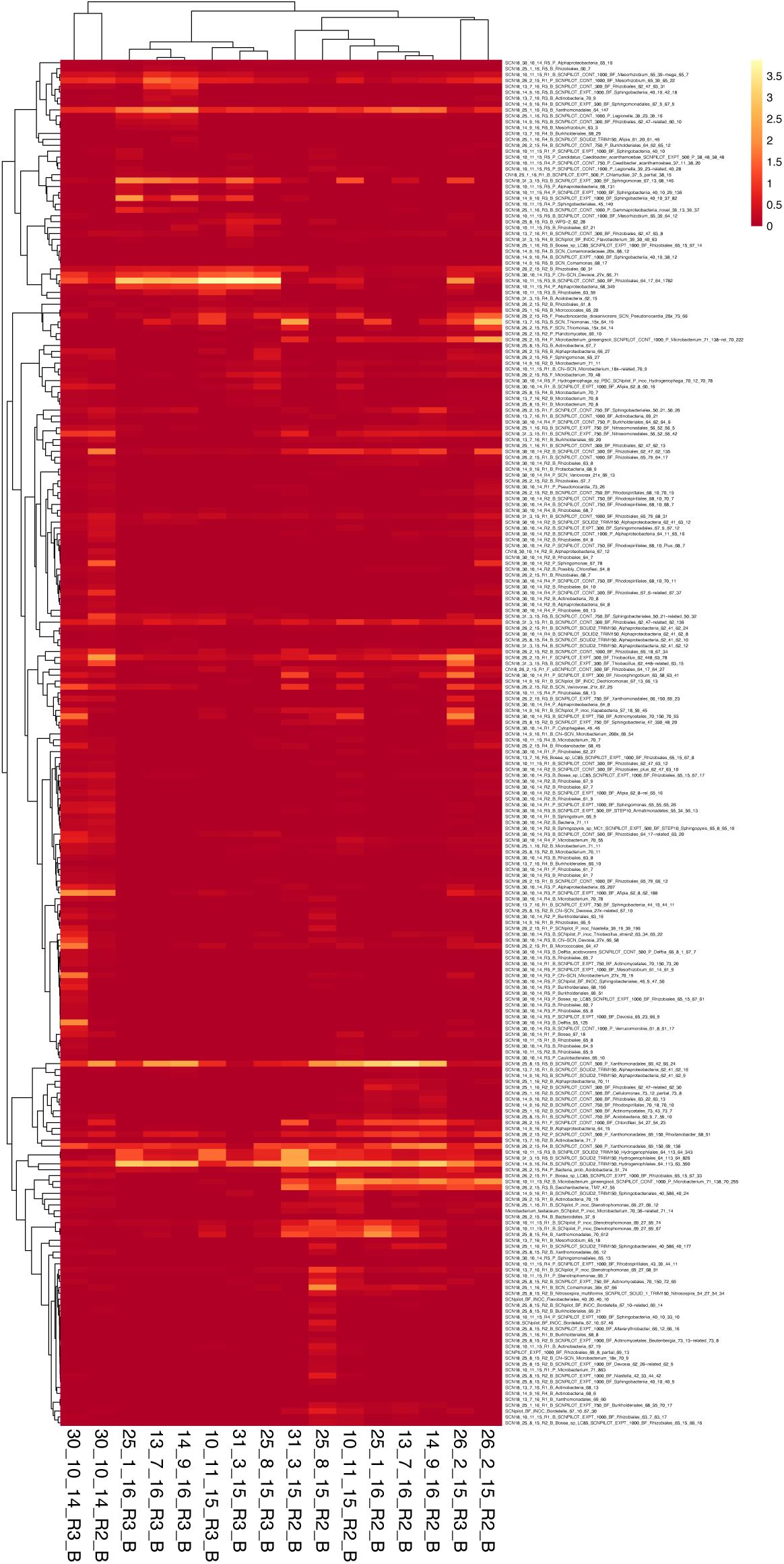
Heatmap based on normalized relative abundance patterns of bacteria in R2 and R3 indicating co-occurrence of Saccharibacteria (TM7) and Microbacterium bins

**Figure S3.**
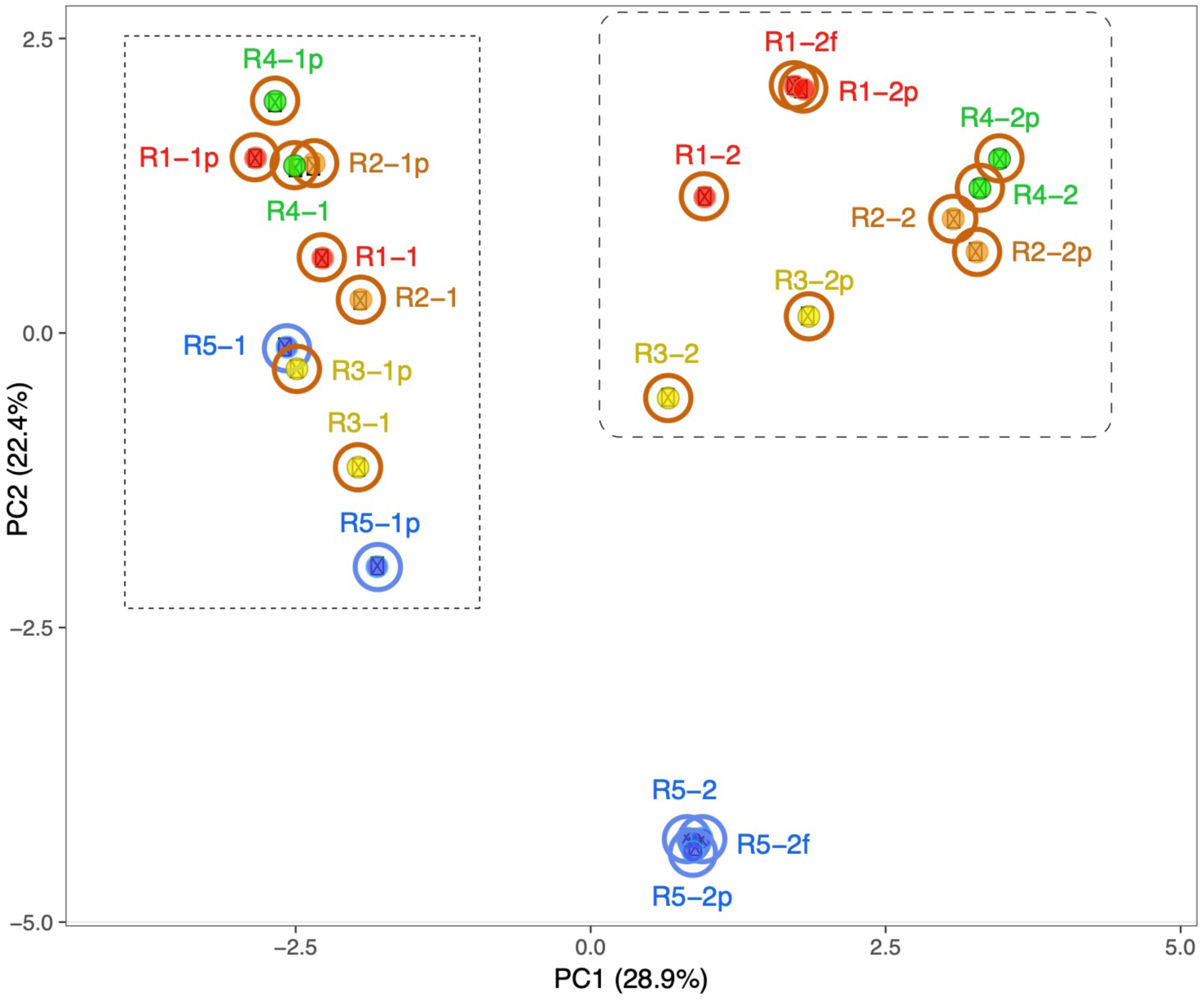
Principal coordinate analysis of community composition at the first two time points in all reactors. The rectangular box shows that communities in all reactors were similar at the first time point, despite reactor 5 receiving no molasses for 117 days. The rounded box indicates communities in all reactors receiving molasses and 250 ppm SCN^-^ (R1, R2, R3 and R4, the second time point). Despite experiencing the same treatments, R1, R2, R3 and R4 are somewhat distinct from each other. Reactor 5 (molasses-free reactor) after the second time point has a very different community composition compared to the other reactors. Included are samples from biofilm, the planktonic fraction (p) and, for reactors 1 and 5, floc samples (f). In most cases, the planktonic fraction is somewhat different from the biofilm (and floc samples).

**Figure S4.**
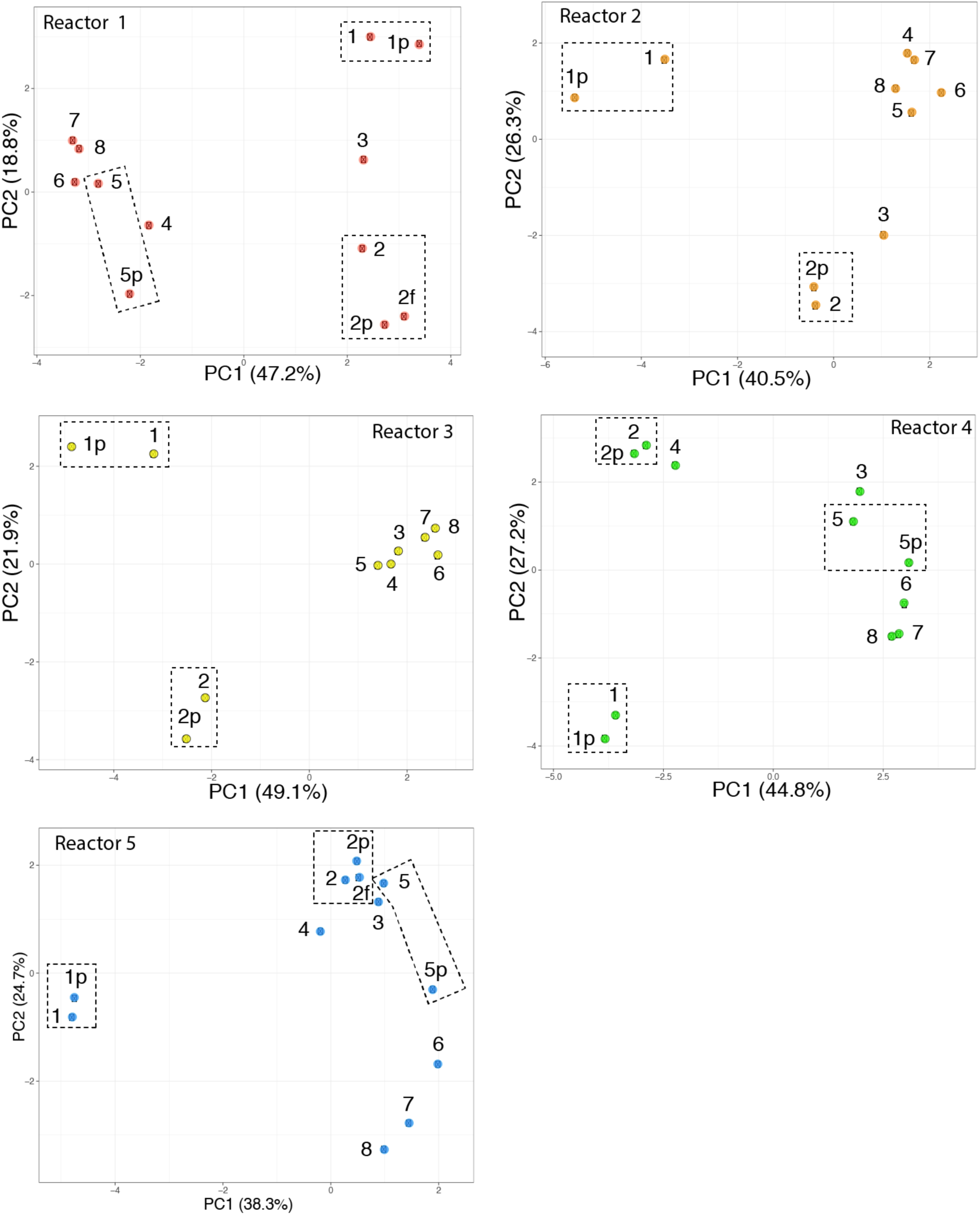
Principal coordinate analysis of community composition for each of the five reactors. Boxes indicate biofilm, planktonic (p) and floc (f) samples from the same time point.

**Figure S5:**
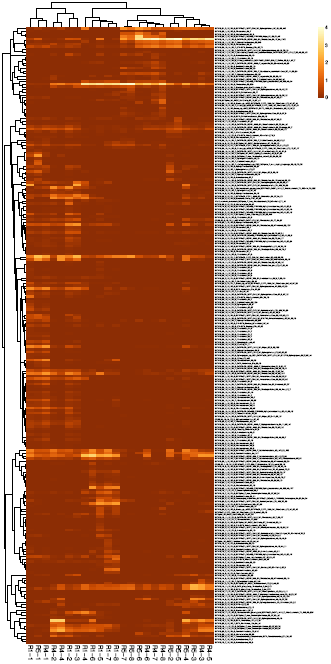
Version of Figure 5B showing organism names

## Supplementary Tables (Excel documents)

**Table S1:** Solution chemistry from the five reactors.

**Table S2**: Percentage of reads from each sample that map to the 232 genomes.

**Table S3**: Abundances of organisms represented by 232 genomes across the samples from the five reactors.

**Table S4:** The relative abundances of bacteria that were abundant in R1 (molasses) and R5 (no molasses) that were amended with thiocyanate and ramped to 2000 mg/L in the five reactors and over time.

## Notes

### Competing Interest Statement

The authors have declared no competing interest.

